# Embryonic expression patterns of panarthropod Teneurin-m/odd Oz genes suggest a possible function in segmentation

**DOI:** 10.1101/762971

**Authors:** Ralf Janssen

**Affiliations:** Uppsala University, Department of Earth Sciences, Palaeobiology, Villavägen 16, Uppsala, Sweden

**Keywords:** odd Oz, odz, pair-rule, segmentation, teneurin, tenascin, Arthropoda, Onychophora, Panarthropoda

## Abstract

**Background:** A hallmark of arthropods is their segmented body, and the so-called *Drosophila* segmentation gene cascade that controls this process serves as one of the best-studied gene regulatory networks. An important group of segmentation genes is represented by the pair-rule genes (PRGs). One of these genes was thought to be the type-II transmembrane protein encoding gene *Tenascin-m* (*Ten-m* (aka *odd Oz*)). *Ten-m*, however, does not have a pair-rule function in *Drosophila*, despite its characteristic PRG-like expression pattern. A recent study in the beetle *Tribolium castaneum* showed that its *Ten-m* gene is not expressed like a segmentation gene, and hence is very unlikely to have a function in segmentation.

**Results:** In this study, I present data from a range of arthropods covering the arthropod tree of life, and an onychophoran, representing a closely related group of segmented animals. At least one ortholog of *Ten-m/odz* in each of these species is expressed in the form of transverse segmental stripes in the ectoderm of forming and newly formed segments – a characteristic of genes involved in segmentation.

**Conclusions:** The new expression data support the idea that *Ten-m* orthologs after all may be involved in panarthropod segmentation.

## Introduction

Teneurins represent a highly-conserved family of type-II transmembrane-protein encoding genes that possess a complex and invariant complement of functional domains (e.g. Tucker, 2018; DePew et al., 2019; Schöneberg and Prömel, 2019). Teneurins are present in at least all of Bilateria (Tucker and Chiquet-Ehrismann, 2006), and although orthologs appear to be absent in sponges, ctenophores and cnidarians, a possible ortholog of bilaterian teneurins has been identified in a choanoflagellate (Tucker et al., 2012). The first teneurin genes were discovered in the vinegar fly *Drosophila melanogaster*. Due to their structural similarity to the extracellular matrix *glycoprotein* Tenascin, these were called *Tenascin-a* (*Ten-a*) (Baumgartner and Chiquet-Ehrismann, 1993) and *Tenascin-m* (*Ten-m*) (Baumgartner et al., 1994), the only two teneurins in *Drosophila*. Interestingly, *Ten-m* was independently identified by another research group and named *odd Oz* (*odz*) after its expression in the brain and the heart (the gifts bestowed by the wizard of Oz) (Levine et al., 1994). In recent years, phylogenetic analyses revealed the presence of both orthologs, *Ten-a* and *Ten-m*, in all arthropods, but also showed that in onychophorans, a closely related outgroup to arthropods, and in another ecdysozoan, the nematode worm *Caenorhabditis elegans*, there is only one teneurin gene (Minet and Chiquet-Ehrismann, 2000; Wides, 2019) (Figure 1A). This gene is similar to both arthropod *Ten-a* and *Ten-m* and thus suggests that the latter are the result of a gene duplication in the stem leading to Arthropoda.

**Figure 1.**
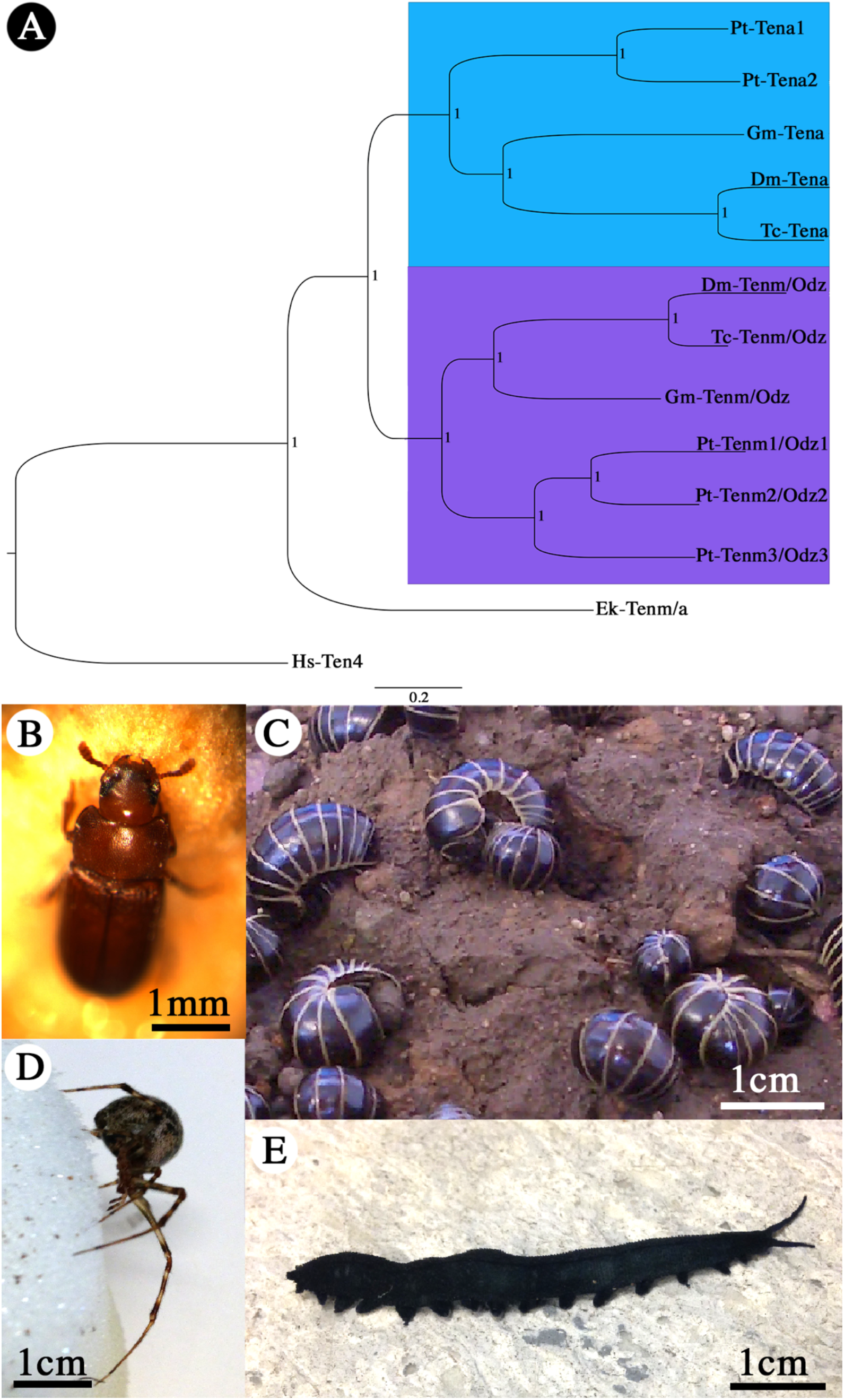
Phylogeny and research organisms. **A** Phylogenetic analysis of teneurin genes. Species abbreviations: Dm, *Drosophila melanogaster* (Hexapoda: Diptera); Ek, *Euperipatoides kanangrensis* (Onychophora); Gm, *Glomeris marginata* (Myriapoda: Diplopoda); Pt, *Parasteatoda tepidariorum* (Chelicerata: Arachnida); Hs, Homo sapiens (Vertebrata); Tc, *Tribolium castaneum* (Hexapoda: Coleoptera); Blue shade: Teneurin-a group. Purple shade: Teneurin-m (Odz) group. Node support is given as posterior probabilities. **B** *Tribolium castaneum*. **C** *Glomeris marginata*. D Adult female of *Parasteatoda tepidariorum*. **E** Adult female of *Euperipatoides kanangrensis*.

Of the two *Drosophila* teneurin genes, *Ten-m* appeared to be the more interesting because it was thought to represent a pair-rule gene (Baumgarnter et al., 1994; Levine et al., 1994), and thus a component of the famous *Drosophila* segmentation gene cascade (StJohnston and Nüsslein-Volhard, 1992). Critical reexamination, however, revealed that *odd paired* (*opa*) and not *Ten-m* was responsible for the reported pair-rule phenotypes, and that *Ten-m* is involved in motor neuron growth and guidance, and not segmentation (Zheng et al., 2011). At some point, a pair-rule function was even attributed to *Ten-a*, but somewhat tragically, even this finding was incorrect (Rakovitsky et al., 2007 (retracted in 2012)). However, since Ten-m protein is expressed in a “typical” pair-rule like pattern in *Drosophila* (Baumgarner and Levine, 1994; Levine et al., 1994), alternative functions in segmentation were discussed, and a possible function as an oscillator in segmentation was suggested (Hunding and Baumgartner, 2017). The very recent work of Jin et al. (2019) investigated the embryonic expression pattern of *Ten-m* in the red flour beetle *Tribolium castaneum*. From their data, Jin and colleagues (2019) concluded that *Ten-m* cannot be involved in segmentation, since it is not expressed in the ectoderm and thus in tissue that is undergoing primary segmentation. The combined data from *Drosophila* and *Tribolium* somewhat intuitively suggest that the pair-rule like expression of *Ten-m* in the highly-derived dipteran fly *Drosophila* may represent a derived feature.

In this paper, however, I present the first comprehensive data on the embryonic expression patterns of teneurin genes (*Ten-m* and *Ten-a* genes) in Panarthropoda, represented by the red flour beetle *Tribolium castaneum* (Hexapoda: Coleoptera), the pill millipede *Glomeris marginata* (Myriapoda: Diplopoda), the common house spider *Parasteatoda tepidariorum* (Chelicerata: Arachnida), and the velvet worm *Euperipatoides kanangrensis* (Onychophora: Peripatopsidae) (Figure 1B-E). The new data show unique expression profiles of each investigated teneurin gene, but also reveal that at least one paralog of *Ten-m* in each species (except for *Tribolium*) is indeed expressed in a pattern that is compatible with a possible function in segmentation. The lack of such expression of *Ten-m* in *Tribolium* may thus represent a derived rather than an ancestral feature of *Ten-m* function.

## Results

### Complement and phylogenetic distribution of panarthropod teneurin genes

Reciprocal BLAST searches revealed the presence of two teneurin genes in *Tribolium* and *Glomeris*, five in the *Parasteatoda* and one in *Euperipatoides* (Figure 1A). The distribution of teneurin genes reflects the relationship of the panarthropod species used in this study (e.g. Campbell et al., 2011) (Figure 1 A). The phylogenetic distribution of these genes suggests that a single teneurin gene was present in the last common ancestor of arthropods and onychophorans, and that this gene was duplicated in the stem leading to the arthropods. In the spider, these genes, *Ten-a* and *Ten-m*, underwent additional duplications. One of these duplications is likely the result of a whole genome duplication that occurred in the last common ancestor of Arachnopulmonata (Schwager et al., 2017) giving rise to *Ten-a1* and *Ten-a2* as well as *Ten-m1/2* and *Ten-m3*. An additional duplication event then lead to the presence of three paralogs of Ten-m in the spider (*Ten-m1/2* duplicating into *Ten-m1* and *Ten-m2*) (Figure 1A).

### Expression of arthropod Teneurin-m (Ten-m) orthologs

*Glomeris Ten-m* is first expressed in the form of transverse segmental stripes in the *regio germinalis* that originates from the early blastoderm (Figure 2A/B) (see Janssen et al., 2004). *Ten-m* is also weakly expressed in the form of a stripe in the segment addition zone (SAZ) (Figure 2A-E). The segmental stripes disappear as the segments mature and resolve into dots of expression in the ventral tissue (likely associated with the developing ventral nervous system), and solid blocks of expression in the dorsal segmental units (Figure 2C-H) (see Janssen, 2011). Additional expression is in the ocular region that harbors the developing brain (Figure 2A-H), the mesoderm of all appendages (Figure 2C-H and Supplementary Figure S1A) and of the anal valves (Figure 2B-H). In late stages, *Ten-m* is also expressed in the developing heart (Figure 2H).

**Figure 2.**
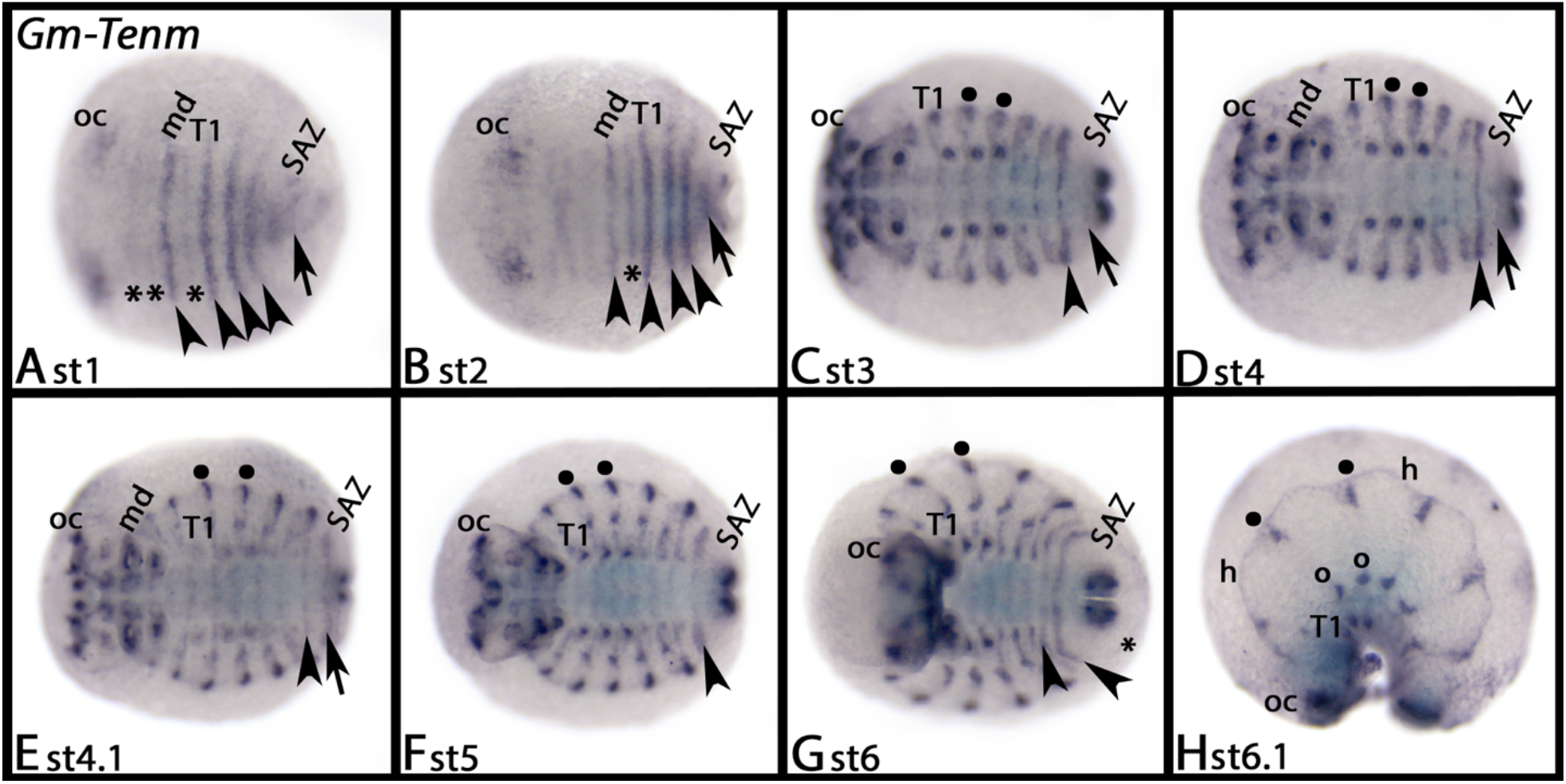
Expression of *Glomeris Ten-m*. In all panels, except for panel H, anterior is to the left, ventral views. Panel H represents a lateral view. Developmental stages are indicated. Arrows point to weak expression in the SAZ. Arrowheads point to transverse stripes of expression. Asterisks in panels A and B mark delayed expression in some segments that originate from the *regio germinalis*. The asterisk in panel G points to expression surrounding the posterior pole of the embryo. Filled circles mark expression in the dorsal segmental units. Open circles in panel H mark lateral domains of expression. Abbreviations: md, mandibular segment; h, heart; oc, ocular region (brain); SAZ, segment addition zone; T1, first walking-leg bearing segment.

*Parasteatoda Ten-m1* is only expressed in late developmental stages when it is in the head lobes, the ventral nervous system, and in the form of a single dot dorsal in the base of the pedipalps and the legs (Supplementary Figure S2).

*Parasteatoda Ten-m2* is expressed in transverse segmental stripes (Figure 3). First, a single broad stripe appears anterior in the forming germ band; this stripe is likely associated with the formation of the labrum and the cheliceral segment (Figure 3A). At subsequent developmental stages, additional stripes form, each associated with a single segment (Figures 3B-H and 4C/D). Later, it becomes clear that the positon of these stripes is located in the forming inter-segmental grooves (Figures 3E, 4C and Supplementary Figure S3). *Parasteatoda Ten-m2* is also expressed in the developing brain in the head lobes (Figures 3B-H and 4A/E/H), the ventral nervous system (Figures 3F/G and 4A-C), and in a complex pattern in the appendages (Figure 4E/F and Supplementary Figure S5B).

**Figure 3.**
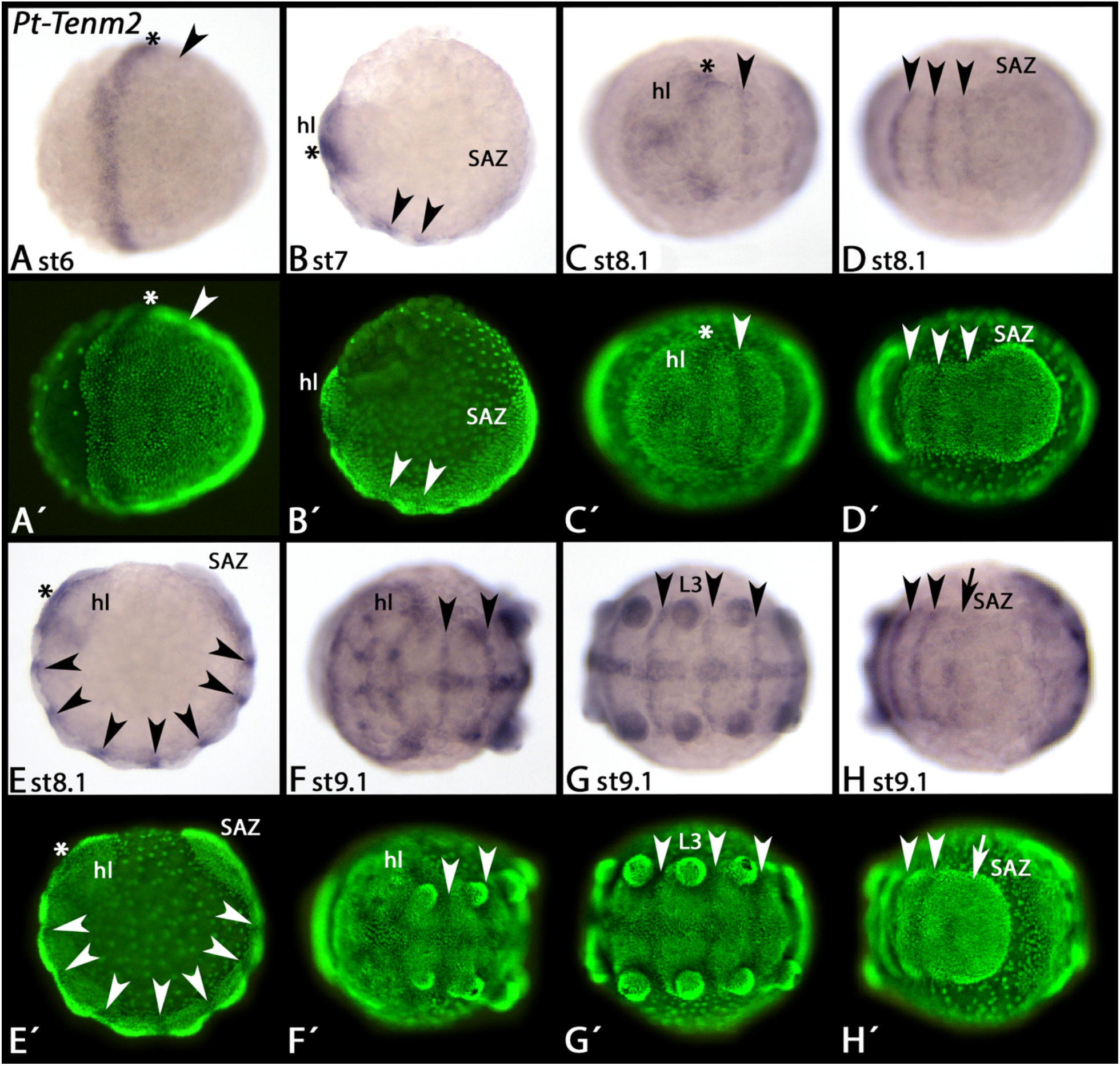
Early expression of *Parasteatoda Ten-m2*. In all panels, anterior is to the left, ventral views (except for panel B, lateral view). Developmental stages are indicated. Panels A’-H’ represent Cybr-Green counter-stained embryos as seen in panels A-H. The arrows in panel H points to weak stripe of expression in the SAZ. Arrowheads mark transverse segmental stripes of expression. Asterisks mark expression in the anterior head (labral and cheliceral segment). Abbreviations: head lobe; L3, third leg-bearing segment; SAZ, segment addition zone.

**Figure 4.**
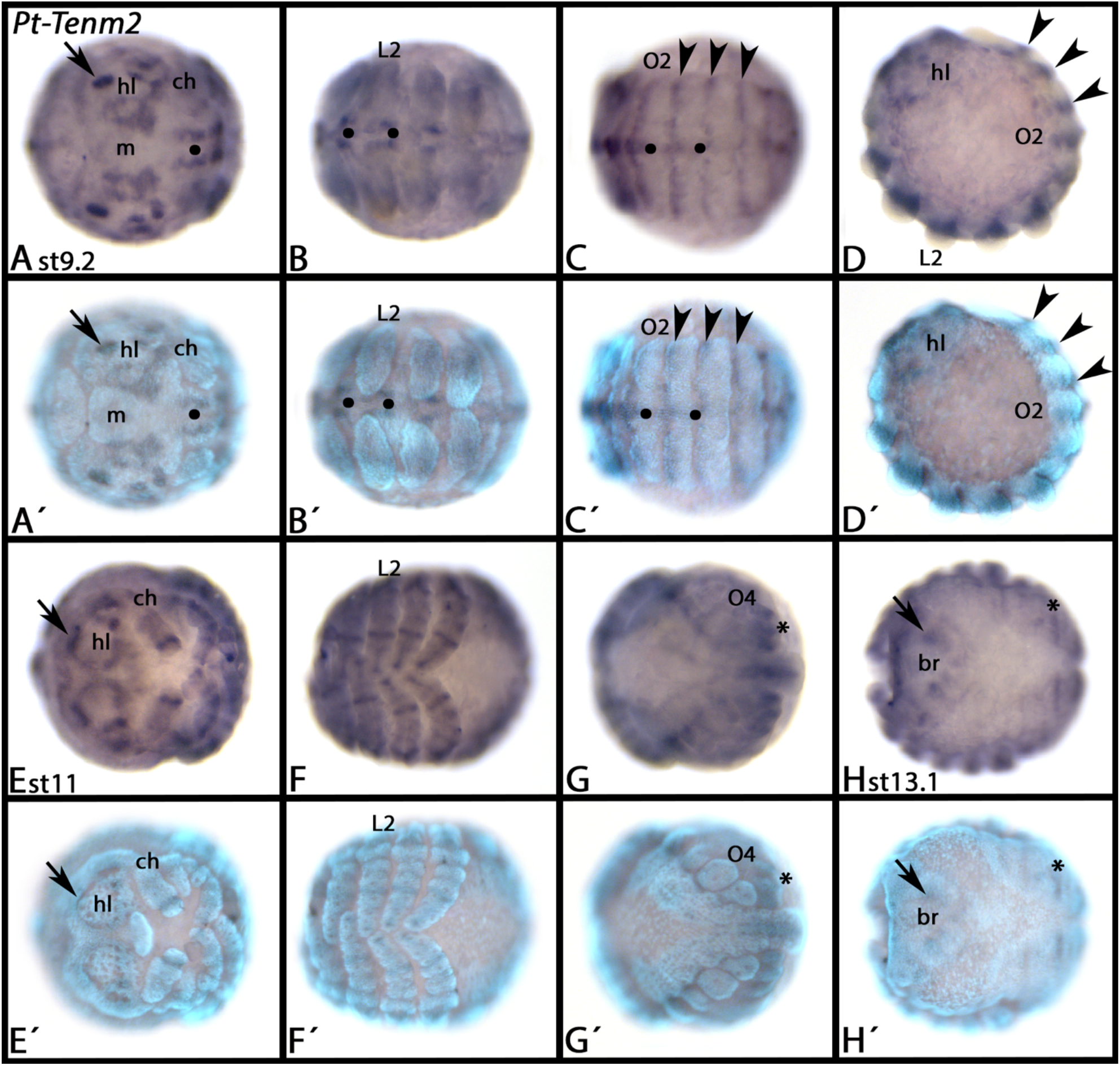
Late expression of *Parasteatoda Ten-m2*. In all panels, anterior is to the left and ventral view (except panel D, lateral view and panel H, dorsal view). Arrowheads point to transverse segmental stripes of expression. Arrows point to expression in the brain. Asterisks in panels G/H mark expression in the alary muscles. Filled circles mark expression in the ventral nervous system. Panels A’-G’ represent Cybr-Green counter-stained embryos as seen in panels A-G. Abbreviations: br, brain; ch, chelicera; hl, head lobe, L2, second walking-leg bearing segment; m, mouth; O2/4, second and fourth opisthosomal segment respectively;

*Parasteatoda Ten-m3* is expressed in dorsal most tissue corresponding to the head lobes (Supplementary Figure S4A/B/E), the prosoma (Supplementary Figure S4A/C) and the opisthosoma (Supplementary Figure S4A-E) with exception of the SAZ (Supplementary Figure S4A/B/D). Later, expression in the opisthosoma likely relates to the development of the alary muscles of the heart (Supplementary Figure S4H/I) (cf. Janssen and Damen, 2008). *Ten-m3* is also expressed in the head lobes (Supplementary Figure S4E/G/I), the mesoderm of the chelicerae and in the form of a distal mesodermal patch in the pedipalps and the legs (Supplementary Figures S4F/J and S1C).

### *Expression of the single onychophoran teneurin gene*, Teneurin-a/m (Ten-a/m)

*Euperipatoides Ten-a/m* is expressed in the frontal appendages (Figure 5), weakly in the mouth and anus (Figure 5A-C), and in the mesoderm of newly formed posterior segments (Figure 5A-D). At later developmental stages, expression appears in the posterior part of the eyes (Figure 5D-F) and the mesoderm of all appendages and the anal valves.

**Figure 5.**
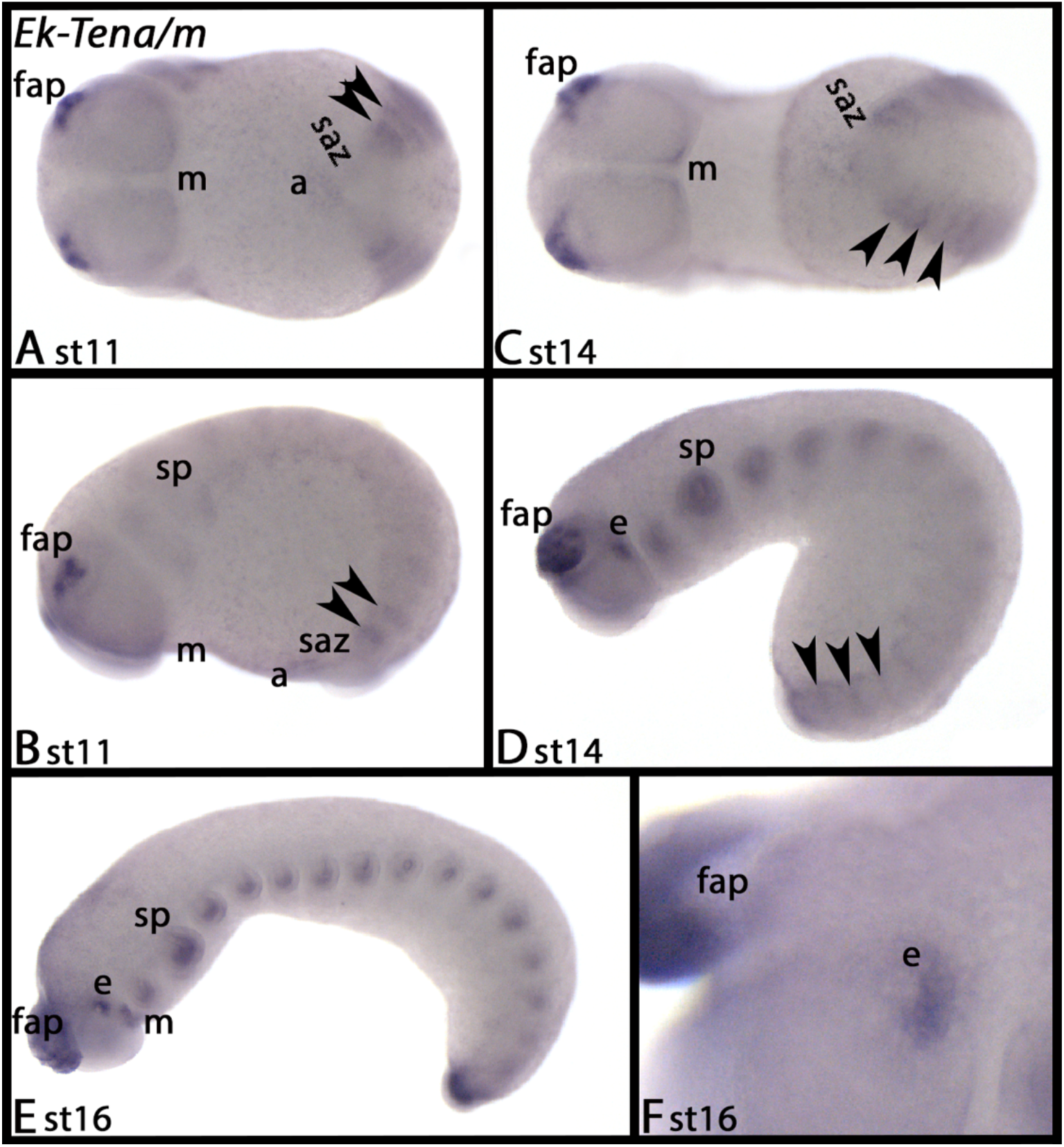
Expression of the single *Euperipatoides* teneurin gene, *Ten-a/m*. In all panels, anterior is to the left. Panels A and B represent ventral views; panels B-F represent lateral views. Arrowheads point to transverse stripes of expression in newly formed segments. Abbreviations: a, anus; e, eye; fap, frontal appendage; m, mouth; saz, segment addition zone; sp, slime papilla.

### Expression of arthropod Teneurin-a (Ten-a) orthologs

*Tribolium Ten-a* is strongly expressed in the developing brain in the ocular region (Supplementary Figure S5A-D), the ventral nervous system (Supplementary Figure S5B-D), and in a subset of cells that likely correspond to the peripheral nervous system (Supplementary Figure S5D). At later stages, it is also expressed in distal region of the developing hindgut (Supplementary Figure S5D).

Like for *Tribolium*, *Glomeris Ten-a* is expressed in the brain and the ventral nervous system, albeit much later and weaker than in *Tribolium* (Supplementary Figure S5E-G). Additional expression is in the form of a small dot at the dorsal base of the antennae and in the anal valves (Supplementary Figure 5E-G).

*Parasteatoda Ten-a1* is expressed in the brain in late developmental stages, the ventral nervous system and the mesoderm of the appendages (except for the labrum) (Supplementary Figures S6A-F and S1D).

*Parasteatoda Ten-a2* is first expressed in the ventral nervous system associated with the prosomal segments (Supplementary Figure S6G-L). In the pedipalps and the legs, but not the other appendages, *Ten-a2* is expressed in the form of a sub-terminal domain (Supplementary Figures S6G/K/L and S1E). At later developmental stages, *Ten-a2* is also expressed in the brain and in the developing book lungs and tracheal lungs in the second and third opisthosomal segment (Supplementary Figure S6L).

## Discussion

### A possible function of Ten-m/odz in arthropod segmentation

The *Drosophila* segmentation gene cascade (SGC) arguably is the most famous and one of the best-investigated gene regulatory network. This hierarchic gene interaction is in control of anterior-posterior body patterning of the fly, and thus the process of segment formation (e.g. Nüsslein-Volhard and Wieschaus, 1980; Ingham, 1988; Cohen and Jürgens, 1991). *Drosophila* development represents a derived form of segmentation in which most of its body is patterned/segmented (almost) simultaneously from a uniform blastoderm (e.g. Pankratz and Jäckle, 1993). Many of the genes involved in this process, however, likely play similar function(s) in other arthropods and their closest relatives, the tardigrades and the onychophorans, all of which add segments sequentially from a posterior segment-addition zone (reviewed in e.g. Peel et al., 2005; Damen, 2007; Smith and Goldstein, 2017; Janssen, 2017; Dunlop and Lamsdell, 2017). One important level of the SGC is represented by the pair-rule genes (PRGs). PRGs are transcription factor-encoding genes that are typically expressed in the form of a unique pattern of seven transverse stripes in the late blastoderm of the fly (e.g. Harding et al., 1986; Carroll and Scott, 1986; Gergen and Butler, 1988; Carroll et al., 1988). In arthropods with sequential addition and patterning of segments, the orthologs of these genes are expressed in the form of transverse stripes or dynamic domains in the ectoderm of the posterior segment addition zone (SAZ) or newly formed segments, or are expressed as transverse stripes in the anterior segments that derive from the blastoderm (e.g. Damen et al., 2005; Choe et al., 2006, 2017; Choe and Brown, 2007; Janssen et al., 2011, 2012; Eriksson et al., 2013; Auman and Chipman, 2018).

In a recent study, Jin and colleagues (2019) demonstrated that in the beetle *Tribolium*, *Ten-m* is not expressed like a segmentation gene, and thus is not involved in segmentation.

In *Glomeris* and *Parasteatoda*, however, at least one ortholog of Ten-m is expressed in the form of transverse segmental stripes in newly forming segments, both in anterior segments that originate from the early blastoderm (or the germ disc in the spider), and in segments that are added from the posterior SAZ (Figures 2–4 and S3). This suggests a possible function in segment formation, or maintenance of segmental boundaries. The expression of *Glomeris Ten-m* and *Parasteatoda Ten-m2* is very similar to that of other segmentation genes including the pair-rule gene orthologs in these species (Damen et al., 2005; Janssen et al., 2008, 2011, 2012; Janssen, 2012; Schönauer et al., 2016; Hemmi et al., 2018). In *Glomeris*, the primary PRGs *runt* (*run*) and *even-skipped* (*eve*) both are expressed in the form of one (or more) circles around the posterior pole (anus) of the developing embryo, representing a unique feature of PRG expression (Janssen et al., 2011); and this detail of PRG expression is also present for *Ten-m* (Figure 2G).

The phylogenetic distribution of segmentation-gene like expression of Teneurin-m genes in Arthropoda thus suggests that the lack of such expression in *Tribolium* is a derived character. The single teneurin gene of the onychophoran *Euperipatoides* is also expressed in transverse segmental stripes in newly formed segments (Figure 5). In onychophorans, however, the PRG-system appears to be little conserved (Janssen and Budd, 2013), and the segmentation-gene like expression of *Ek-Ten-a/m* is not as obvious as in arthropods. Notably, however, the expression of *Ten-a/m* is very similar to that of the only PRG ortholog that is possibly involved in onychophoran segmentation (or at least posterior elongation), *even-skipped* (*eve*) (Janssen and Budd, 2013). Thus, the expression of *Ten-a/m* could indicate that a possible segmentation-gene (or related) function may date back to the last common ancestor of Panarthropoda, and that such function was already established before the duplication of an ancestral panarthropod teneurin gene (*Ten-a/m*) into *Ten-m* and *Ten-a*. It was therefore necessary to investigate whether such function/expression could have been retained in the *Ten-a* orthologs of arthropods including the beetle *Tribolium* that does not express *Ten-m* in a segmentation-gene like pattern (Jin et al., 2019). Gene expression analysis of *Ten-a* orthologs in all investigated arthropods, including *Tribolium*, however, revealed that neither of them is expressed like a potential segmentation gene, revealing that the segmentation-gene like expression of arthropod teneurin genes is a feature of *Ten-m*, and that the lack of such expression in *Tribolium* indeed likely represents a derived character.

Further investigation including functional studies may reveal if arthropod *Ten-m* genes indeed are involved in segmentation, and what their function(s) in this process may be.

## Experimental Procedures

### Research organisms and embryos

Research animals and their embryos were handled as described in Grossmann and Prpic (2012) (*Tribolium castaneum*), Janssen et al. (2004) (*Glomeris marginata*), Prpic et al. (2008) (*Parasteatoda tepidariorum*), and Hogvall et al. (2014) (*Euperipatoides kanangrensis*). Developmental staging follows Strobl and Stelzer (2014) (*Tribolium*), Janssen et al. (2004) (*Glomeris*), Mittmann and Wolff (2012) (*Parasteatoda*), and Janssen and Budd (2013) (*Euperipatoides*).

### *RNA extraction, gene cloning, whole mount* in-situ *hybridization, and nuclear counter staining*

Total RNA was extracted with TRIZOL (Invitrogen) from a mix of embryos representing all stages from the blastoderm stage to hatching. Total RNA was then reverse transcribed into cDNA using the SuperScript IV Reverse Transcriptase (Invitrogen). Gene fragments were amplified by RT-PCR with gene-specific primers based on sequenced genomes (*Tribolium* and *Parasteatoda*) and sequenced embryonic transcriptomes (*Glomeris* (SRA accession: PRJNA525752) and *Euperipatoides* (SRA accession: PRJNA525753)). In all cases, a nested PCRs was performed, using an initial PCR as template. Primer sequences are listed in Supplementary Table 1. Gene fragments were cloned into the PCRII vector (Invitrogen) and sequenced on an ABI3730XL automatic sequencer (Macrogen, Seoul, South Korea). Unique gene identifiers are listed in Supplementary Table 2. *In-situ* hybridization was performed as described in Janssen et al. (2018) using BM Purple (Roche) as staining substrate. For confocal microscopy, embryos were stained with SIGMAFAST Fast Red TR/NaphtolAS-MX (SIGMA). Morphology of the embryos was visualized with the nuclear dye SYBR Green in phosphate buffered saline with 0.1% Tween-20 (PBST-0.1%).

### Phylogenetic analysis

Reciprocal BLAST searches applying tblastn, blastp and blasty were run with the *Drosophila melanogaster* sequences of Tenascin-m (aka Odd Oz (Odz)) and Tenascin-a to identify teneurin genes. Amino acid sequences were aligned using T-Coffee with default parameters in MacVector v12.6.0 (MacVector, Inc., Cary, NC), or Aliview 1.18.1 for Linux (Larsson, 2014). The phylogenetic analysis was performed with MrBayes (Huelsenbeck and Ronquist, 2001) with a fixed WAG amino acid substitution model with gamma-distributed rate variation across sites (with four rate categories), unconstrained exponential prior probability distribution on branch lengths, and exponential prior for the gamma shape parameters for among-site rate variation. The phylogenetic tree was calculated applying 300000 cycles for the Metropolis-Coupled Markov Chain Monte Carlo (MCMCMC) analysis (four chains; chain-heating temperature of 0.2). Markov chains were sampled every 200 cycles. 25% of samples were applied as burn-in. Clade support was determined with posterior probabilities in MrBayes. Unique sequence identifiers for all sequences used in the phylogenetic analysis are listed in Supplementary Table 2.

### Data documentation

A Leica DC490 digital camera equipped with a UV light source mounted onto a MZ-FLIII Leica dissection microscope was used for documentation of stained embryos. Confocal microscopy was performed using an inverted Leica TCS SP5 confocal microscope. For the detection of Fast Red and DAPI signal. Emission wavelengths for Fast Red and DAPI were 561nm and 404nm, respectively. Collected wavelengths for Fast Red were between 600nm and 642nm, and for DAPI were between 430nm and 550nm. Whenever appropriate, contrast and brightness were adjusted with the image-processing software Adobe Photoshop CC2018 for Apple Macintosh (Adobe Systems Inc.).

## Acknowledgments

I would like to acknowledge the support of the New South Wales Government Department of Environment and Climate Change by provision of a permit SL100159 to collect onychophorans at Kanangra-Boyd National Park, and Glenn Brock, David Mathieson, Robyn Stutchbury and Noel Tait for their help during onychophoran collection. Embryos of *Parasteatoda* and *Tribolium* were kindly provided by Matthias Pechmann, Anna Schönauer, Alistair McGregor and Gregor Bucher. The picture of an adult *Tribolium castaneum* shown in Figure 1 is a gift from Gregor Bucher.

