## Supplementary Data for "Embryonic expression patterns of panarthropod Teneurin-m/odd Oz genes suggest a possible function in segmentation"

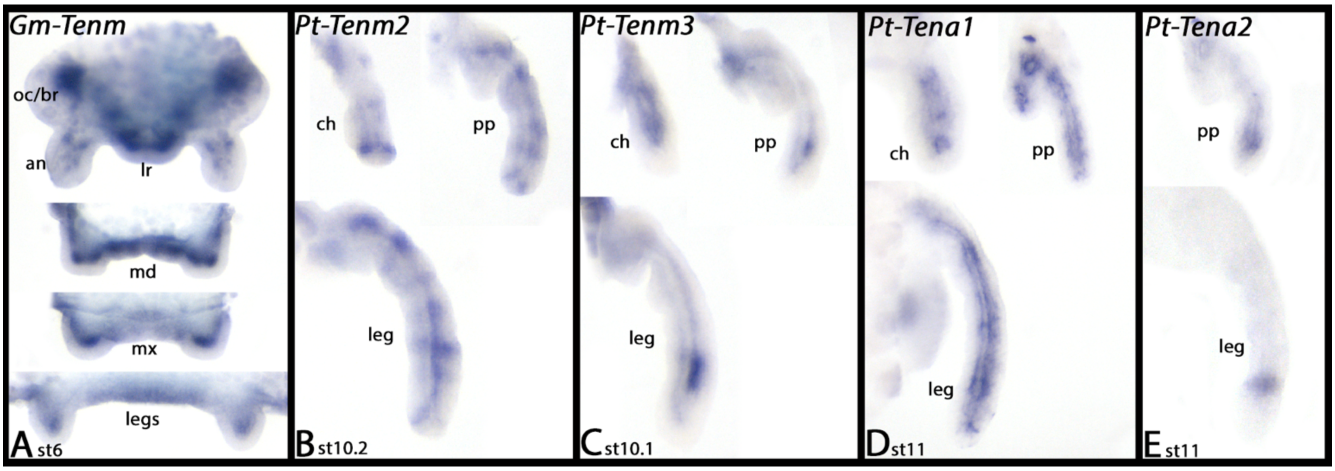

Supplementary Figure 1 – Appendicular expression of teneurin genes

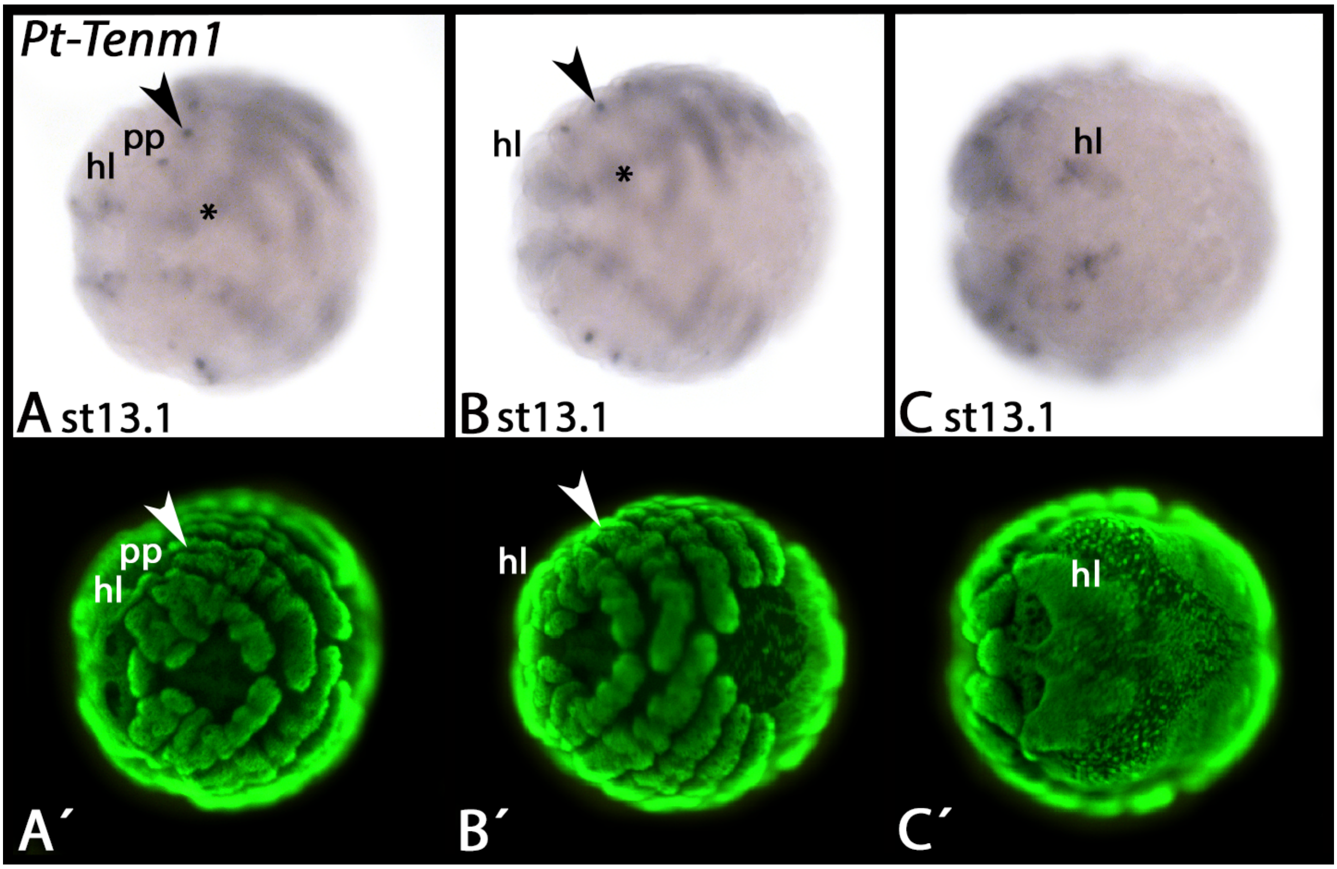

Supplementary Figure 2 – Expression of *Parasteatoda Ten-m1*

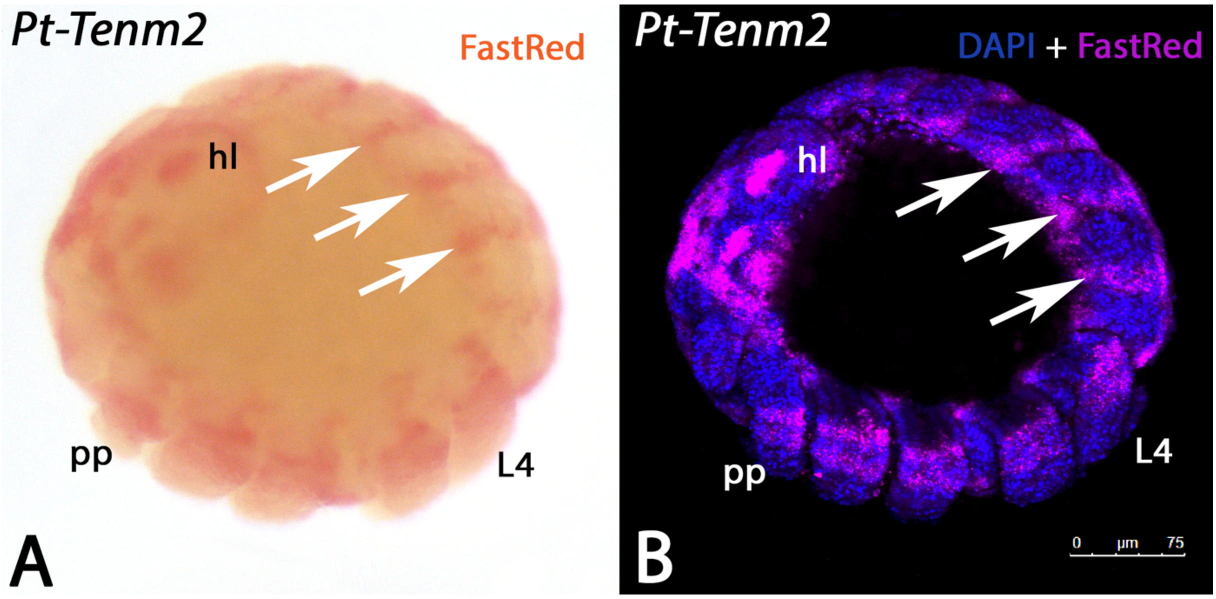

Supplementary Figure 3 – FastRed staining and confocal microscopy of *Parasteatoda Ten-m2*

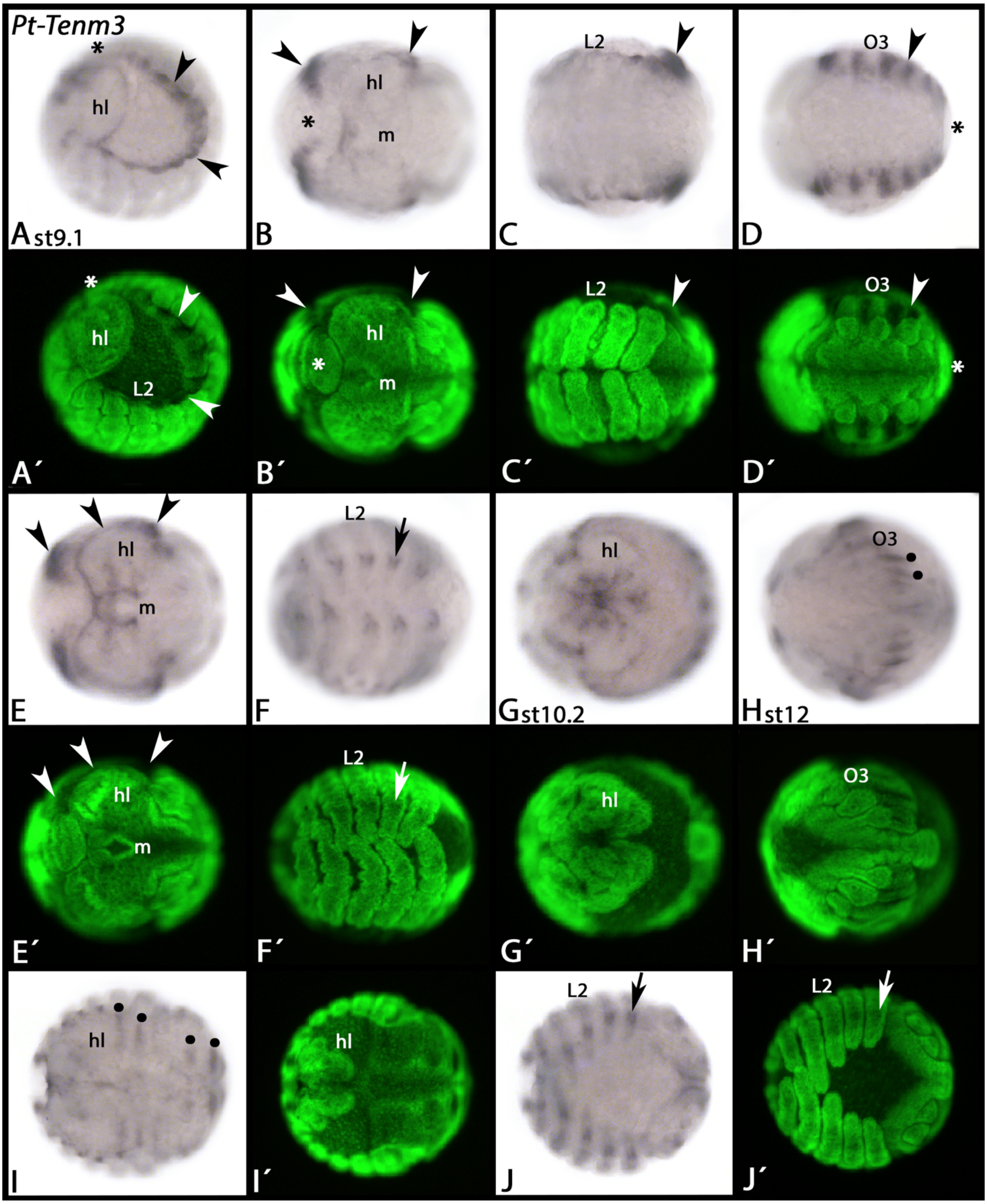

Supplementary Figure 4 – Expression of *Parasteatoda Ten-m3*

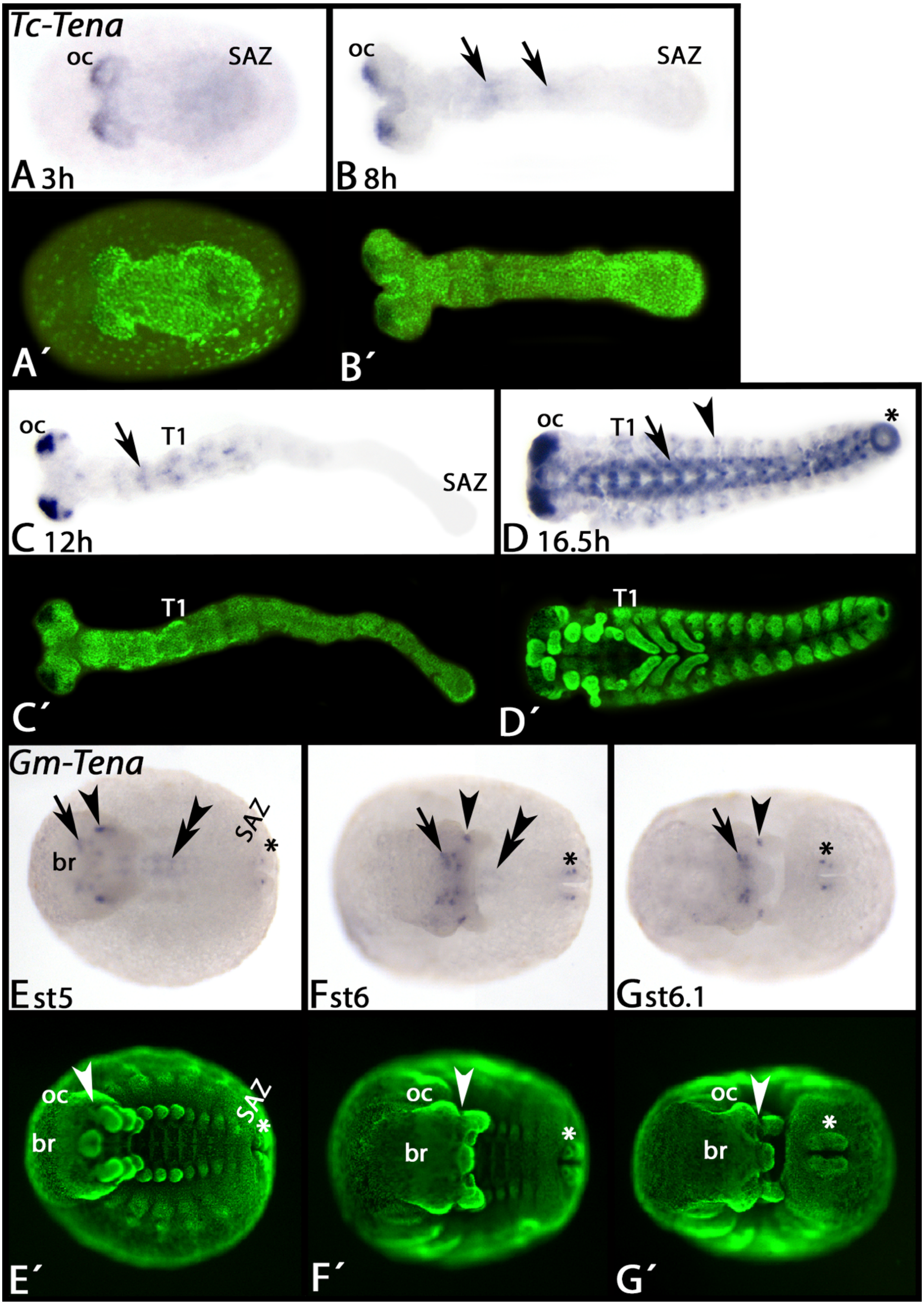

Supplementary Figure 5 – Expression of *Tribolium Ten-a* and *Glomeris Ten-a*

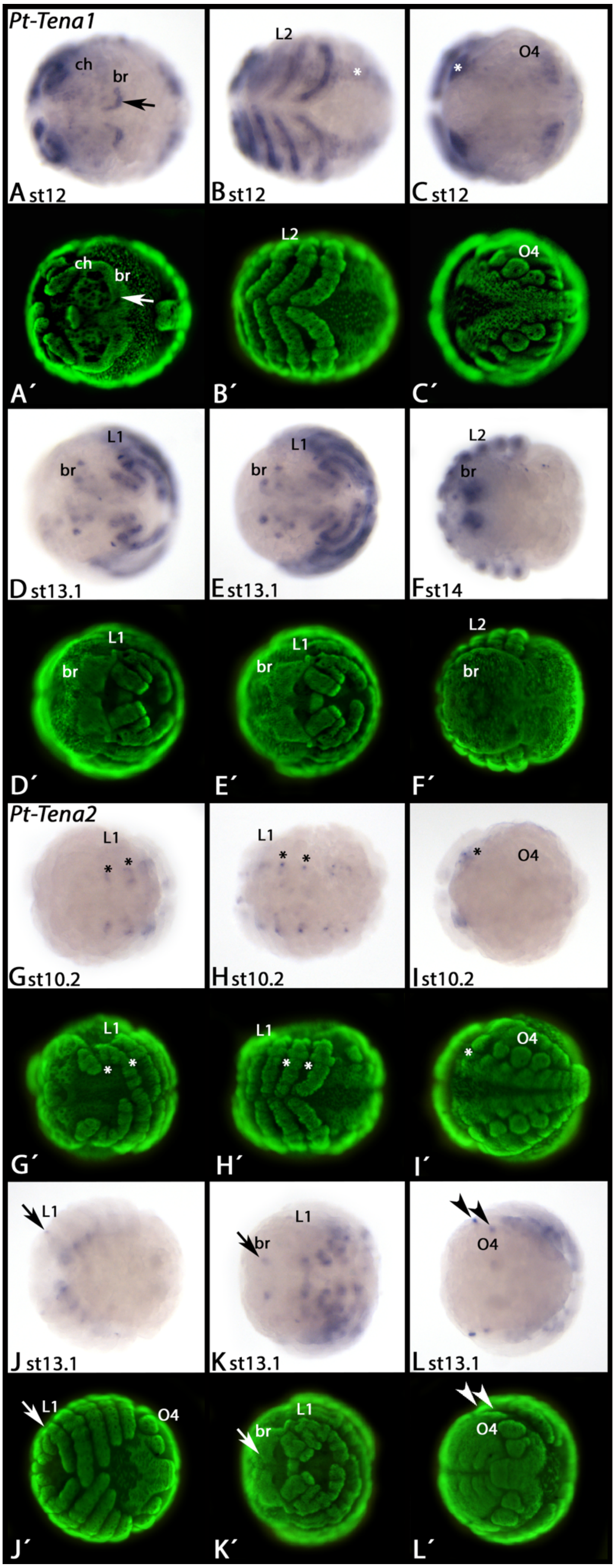

Supplementary Figure 6 – Expression of *Parasteatoda Ten-a1 and Ten-a2*

Supplementary Figure S1 – Appendicular expression of teneurin genes

A Mesodermal expression in all myriapod appendages. B Ectodermal rings of expression. C Mesodermal expression in the chelicerae and in the form of a distinct distal mesodermal patch in the pedipalps and the legs. D Mesodermal expression in chelicerae, pedipalps and legs. E Expression in the form of a sub-distal domain in the pedipalps and the legs, but not the chelicerae (not shown). Abbreviations: br, brain; ch, chelicera; lr, labrum; md, mandible; mx, maxilla; oc, ocular region; pp, pedipalp.

Supplementary Figure S2 – Expression of *Parasteatoda Ten-m1*

In all panels, anterior is to the left. Panel A, anterior view. Panel B, ventral view. Panel C, dorsal view. Panels A´-C´ represent Cybr-Green counter-stained embryos as shown in panels A-C. Arrowheads point to expression at the base of the prosomal appendages. Abbreviations: hl, head lobe; pp, pedipalp.

Supplementary Figure S3 – FastRed staining and confocal microscopy of *Parasteatoda Ten-m2*

A Bright field photography of a FastRed-stained stage 9.2 embryo. B Same embryo as shown in panel A in similar position. Confocal microscopy reveals expression of *Ten-m2* (FastRed, magenta signal) in the form of ectodermal segmental stripes (arrows). The blue signal represents DAPI-staining of nuclei. Abbreviations: hl, head lobe; L4, fourth walking-leg; pp, pedipalp

Supplementary Figure S4 – Expression of *Parasteatoda Ten-m3*

All panels, anterior to the left and ventral views (except for panel A, lateral view; panel I, dorsal view). Panels A´-J´ represent Cybr-Green counter-stained embryos as seen in panels A-J. Arrowheads point to expression along the dorsal margin of the embryos. Arrows point to expression around the limbs. Filled circles point to expression in the alary muscles. The asterisks mark the SAZ that does not express *Ten-m3*. Abbreviations: hl, head lobe; L2, second walking-leg bearing segment; m, mouth; O3, third opisthosomal segment.

Supplementary Figure S5 – Expression of *Tribolium* and *Glomeris Ten-a*

In all panels, anterior is to the left, ventral views. Panels A´-G´ represent Cybr-Green counter-stained embryos as shown in panels A-G. Arrows in panels B-D point to expression in the ventral nervous system. The arrowhead in panel D points to expression in the peripheral nervous system. The asterisk in panel D marks expression in the hindgut. In panels E-G, arrows mark expression in the brain, single arrowheads point to a distinct domain posterior to the antennae, and double-arrowheads mark expression in the ventral nervous system. The asterisks in panels E-G mark expression in the anal valves. Abbreviations: br, brain; oc, ocular region; SAZ, segment addition zone; T1, first thoracic segment.

Supplementary Figure S6 – Expression of *Parasteatoda Ten-a1* and *Ten-a2*

In all panels, anterior is to the left and ventral views (except panels D, E and K, anterior view; panel F, dorsal view, and panel L, posterior view). Panels A´-L´ represent Cybr-Green counter-stained embryos as shown in panels A-L. Arrows point to expression in the brain. Arrowheads in panel L point to expression in the second and third opisthosomal segment in the primordia of the book lungs and tracheal lungs. Asterisks mark expression in the ventral nervous system.

Abbreviations: br, brain; ch, chelicera; L1/2, first and second walking-leg bearing segment respectively; O4, fourth opisthosomal segment.

| | |
|---|---|
| EkTenM/A_fw1 | TCTGAGGAGGGGATATTGA |
| EkTenM/A_bw2 | TCACCAAACCAGCAAGAGAG |
| EkTenM/A_bw1 | GTGTTGATAGTAACTGGGGC |
| EkTenM/A_fw2 | GCTAGTGACAGGCTCGTTAA |
| TcTenA_fw1 | TCCAAGAAGGGTGCGAAACA |
| TcTenA_bw2 | TCCGCACTATTCACCGTACT |
| TcTenA_bw1 | ATGTCGCATTCGGCACCTTT |
| TcTenA_fw2 | AGAACGGGGCATACACTCTT |
| GmTenA_fw1 | CAGAGAGCAAGGATGGTTTG |
| GmTenA_bw2 | GGAACTCCCATCAGAATCTC |
| GmTenA_bw1 | TCTTAGCCTTCTGTTGTGCC |
| GmTenA_fw2 | AGAAAAGTTCGCCATCCCAG |
| PtTenA1_fw1 | GTGAATGTGAATCTGAGCCG |
| PtTenA1_bw2 | CAATGTTGACAGCATACCCG |
| PtTenA1_bw1 | CACCATCTACGCATTTCCCT |
| PtTenA1_fw2 | GCAAGTTCACAATCCGTTGC |
| PtTenA2_fw1 | TCGGCTCAATAGGCAAGGTT |
| PtTenA2_bw2 | CAGCAGTTCGGATAATGAGG |
| PtTenA2_bw1 | AAATGGAACCTGGACGAAGC |
| PtTenA2_fw2 | CCGAAAATGCTGAGGTTGCT |
| GmTenM_fw1 | CCCTTGCTGTCTCCATATCA |
| GmTenM_bw2 | TGACATTCTTCACCTGCGAG |
| GmTenM_bw1 | CACTATCCTCGCAATCAACC |
| GmTenM_fw2 | GATACCACAGTCAAGACCGT |
| PtTenM1_fw1 | CCAGCAAAGGGAATCACAGT |
| PtTenM1_bw2 | ATACCCTGTCCAGTGAACCT |
| PtTenM1_bw1 | ATCCATTGCTCATCACAGGC |
| PtTenM1_fw2 | TTGTCCGTGTCCTGAGCAAT |
| PtTenM2_fw1 | ACTGCTGATGAATCCGACTC |
| PtTenM2_bw2 | CGAACGCCAAACGACATTAC |
| PtTenM2_bw1 | CGGTGGAGGTGTTTTCTTTG |
| PtTenM2_fw2 | TCATCTACCATCACAGGGCA |
| PtTenM3_fw1 | TTCCGTCCACAATCCCACAA |
| PtTenM3_bw2 | CGAGCAGAAGAACAAAACGC |
| PtTenM3_bw1 | GGAGATTCTTCTGCAATGGC |
| PtTenM3_fw2 | CAATCATCACAGGACAAGGG |

Supplementary Table 1 – Primers

| | |
|---|---|
| Gm-TenM | c59480_g1 |
| Gm-TenA | c57511_g2 |
| Ek-TenM/A | c2122770_g1 |
| Tc-TenA | XP_015834850.1 |
| Pt-TenA1 | XP_015923788.1 |
| Pt-TenA2 | xP_015906484.1 |
| Pt-TenM1 | XP_015928268.1 |
| Pt-TenM2 | XP_021000186.1 |
| Pt-TenM3 | XP_021002483.1 |
| Dm-TenA | NP_001138189.1 |
| Dm-TenM/odz | AAB88281.1 |
| Tc-TenM/odz | XP_015834526 |
| Hs-Ten4 | NP_001092286.2 |

Supplementary Table 2 – Accession Numbers
